# Transcriptomic signature of Human Cardiac Fibroblast in Hypertrophic Obstructive Cardiomyopathy

**DOI:** 10.64898/2026.07.12.738095

**Authors:** Ayman M. Ibrahim, Sarah Halawa, Lydia Sidarous, Aya Galal, Sara Elshorbagy, Hasnaa A. Elfawy, Manar Osman, Faisal Mohamed, Mohamed Roshdy, Mohamed Hosny, Arwa Kohela, Mona Allouba, Ahmed Mostafa, Yasmine Aguib, Magdi Yacoub

**Affiliations:** Faculty of Health and Medical Sciences, University of Surrey, Guildford, GU27AL, UK; Department of Zoology, Faculty of Science, Cairo University, Giza 12C13, Egypt; Aswan Heart Center, Magdi Yacoub Heart Foundation, Aswan, Egypt; Systems Genomics Laboratory, American University in Cairo, New Cairo, Egypt; Department of Biology, American University in Cairo, New Cairo, Egypt; Cardiology Department, Faculty of Medicine, Cairo University, Giza 115C2, Egypt; National Heart and Lung Institute, Imperial College London, London SW3 CLY, UK

**Keywords:** Cardiac Fibroblasts, Hypertrophic Obstructive Cardiomyopathy, Myocardial Fibrosis, Extracellular Matrix, Inflammation

## Abstract

**Background:** Hypertrophic Cardiomyopathy (HCM) is a cardiac disorder characterized by an increased interstitial fibrosis and Extracellular Matrix (ECM) remodeling. Cardiac fibroblasts (CFs) have a crucial role in ECM remodeling as well as influencing contractility. There is accumulating evidence that the phenotype of CFs is disease-specific. Here, we investigate the transcriptome signature of CFs in Hypertrophic obstructive Cardiomyopathy (HOCM) patients and its functional significance.

**Methods:** Primary CFs were isolated from myectomy specimens of 12 clinically phenotyped HOCM patients and 3 controls. RNA libraries were prepared from cell isolates for whole transcriptome sequencing. Differential expression analysis was conducted using the Tuxedo pipeline. Pathway and Gene Ontology (GO) enrichment were performed using Protein-protein interaction network analysis and functional clustering were performed using Cytoscape StringApp. Expression of candidate genes and proteins was validated using quantitative PCR, Immunocytochemistry, immunohistochemistry, cytokines/chemokines profiling and western blotting, in patient-derived CFs extracts, CFs-conditioned media, and myocardial tissue sections.

**Results:** Whole transcriptome analysis identified 265 significant differentially expressed genes (DEGs) in HOCM fibroblasts compared to controls. The most significant GO terms identified were associated with ECM organization and inflammatory response. The most significant GO terms identified were associated with ECM organization and inflammatory response, with circos plot analysis further highlighting pathway-specific gene overlap within inflammatory and structural signaling clusters. The DEGs encompassed gene families such as collagens, proteases, fibulins, inflammatory cytokines, integrins and signaling receptors and kinases. *MYC* was upregulated alongside chemokine ligands and receptors, highlighting a MYC-linked chemokine signaling axis within the inflammatory HOCM-CFs phenotype. The protein expression of selected ‘extracellular Matrix organization’ and ‘inflammatory response’ genes confirmed the transcriptome results.

**Conclusion:** Transcriptomic profiling of patient-derived HOCM-CFs identified genes and pathways associated with inflammatory signaling, ECM remodeling, and altered cell–cell/matrix communication. These findings show that advanced HOCM-CFs acquire an inflammatory-remodeling phenotype, with ECM genes other than collagens.

**Novelty and Significance:** *What Is Known?:*

- Hypertrophic cardiomyopathy is characterized by myocardial hypertrophy, interstitial fibrosis, extracellular matrix remodelling, and inflammatory signalling.
- Most human HCM transcriptomic studies have been performed using whole myocardial tissue, which limits resolution of fibroblast-specific disease programmes.

*What New Information Does This Article Contribute?:*

- Patient-derived cardiac fibroblasts from patients with advanced hypertrophic obstructive cardiomyopathy exhibit a distinct transcriptomic signature enriched for inflammatory signalling, extracellular matrix organization, chemokine activity, and altered cell–matrix communication.
- The HOCM-CF signature is not characterized by uniform collagen gene induction, but by selective ECM remodelling involving fibulins, proteases, integrins, matricellular genes, and downregulation of selected collagen-associated matrix components.
- Comparison with bulk myocardial RNA-sequencing and publicly available human cardiomyopathy data supports the biological relevance of the fibroblast-derived signature while highlighting the limited sensitivity of bulk tissue transcriptomics for resolving fibroblast-specific programmes.

*What Is the Significance?:* This study identifies a disease-associated inflammatory and ECM-remodelling state in patient-derived HOCM cardiac fibroblasts. The findings extend the role of cardiac fibroblasts in HOCM beyond classical collagen deposition and suggest that fibroblast-mediated matrix remodelling, cytokine/chemokine signalling, and altered cell–matrix communication may contribute to myocardial remodelling in advanced obstructive disease. These data support the value of fibroblast-focused profiling for uncovering disease-relevant mechanisms that may be masked in whole myocardial tissue analyses and provide candidate pathways for future mechanistic and therapeutic studies.

## Introduction

Hypertrophic cardiomyopathy (HCM) is an inherited cardiac disorder, characterized by left ventricular thickening, and Extracellular Matrix (ECM) remodeling, generally represented in an increased interstitial fibrosis ^1,2^. Clinical and pathological presentations vary among patients which makes it a pleiotropic disease with different responses to treatment ^3,4^. Although the disease was thought to be a disease of the sarcomere, there is now a growing realization that other cell types, such as cardiac fibroblasts (CFs), play a major role, preceding sarcomeric dysfunction ^4^.

CFs represent the major non-cardiac cell lineage that maintain the myocardial homeostasis and ECM turnover ^5^. During myocardial injury, CFs become activated and may differentiate to myofibroblasts, with an expression of alpha smooth muscle actin (α-SMA), increased secretion of inflammatory mediators, and increased deposition of ECM proteins, which all represent a stress response that aggravate heart diseases ^6,7^. ECM is an essential component of the myocardial tissue that generally contributes to the ventricular size and shape responsiveness during contractions and imparts primary cellular processes for tissue homeostasis ^8^.

The crosstalk between CFs and cardiomyocytes (CMs), as well as with other non-cardiac cells and their surrounding ECM, is crucial for tissue homeostasis ^9,10^. In pathological conditions, however, all these elements are influenced by altered signaling cascades, leading to a microenvironment that chronically affects CFs phenotype and relevant response ^7,11^.

In HCM, collagens, fibronectin, matrix metalloproteinases, inflammatory cytokines, and other ECM components have been identified as part of the matrix turnover machinery during interstitial fibrosis and hypertrophy ^12^. However, the exact role of CFs and other ECM proteins has not been adequately studied.

In this study, we aimed to investigate the transcriptome signature of CFs in patients with Hypertrophic Obstructive Cardiomyopathy (HOCM), pathological ECM remodeling machinery and the major drivers of disease development.

## Materials and Methods

### Cardiac Fibroblasts isolation

Myectomy specimens were collected from the surgery into phosphate buffer saline (PBS) w/o Ca, Mg, and transported to the cell culture laboratory where the fibroblast isolation was performed immediately. Fibroblasts were isolated using the explant method of isolation - with differential centrifugation, as previously described ^13–15^. CFs isolates were characterized using immunocytochemistry and Flow cytometry.

### RNA extraction and sequencing

Upon confluency in a well of a 6-well plate, RNA was extracted from fibroblasts (Qiagen), RNA yield and quality was measured using Nanodrop (Thermo Scientific) and further analyzed for quality control and RNA integrity with Agilent bioanalyzer (Agilent). Isolated total RNA was sequenced using TruSeq Stranded Total RNA with Ribo Zero Gold kit 1 (Illumina). Libraries were quality controlled for size distribution and yield using Tapestation 4200 with high sensitivity dsDNA assay (Agilent Technologies) and sequenced as 151 bp pair-end reads on 4 lanes of a NextSeq 550 (Illumina) using high-output cartridges. Quality assessment of the paired-end raw reads was done using FastQC ^16^(version 0.11.5) before and after trimming with Trimmomatic ^17^(version 0.36).

### Read alignment, mapping, and differential expression identification

Trimmed reads were aligned to the GENCODE ^18^ GRCh38 genome build using STAR^19^ (version 2.7.10). The Tuxedo protocol^20^ (version 2.2.1) was used for differential expression analysis: assembly was done using Cufflinks, sample annotations and GENCODE annotation were merged with Cuffmerge, quantification of transcripts and genes was performed using Cuffquant and differential expression was performed using Cuffdiff. Differentially expressed genes (DEGs) were filtered using q-value ≤ 0.05 and log fold-change ≥ <u>|1.5|</u>.

### Differential Expression Visualization, Network, and Functional Enrichment Analysis

The R packages CummerBund and ggplot2 ^21^ were used for differential expression visualization. Gene Ontology (GO) and pathway enrichment analysis of the differentially expressed genes was performed with the ToppGene suite ^22^, interrogating many pathway databases including the Reactome Pathway Database. Circos plots of relevant enriched pathways were drawn using the circlize^23^ R package. The protein-protein interaction network was constructed using the Cytoscape^24^ (version 3.10.1) with the StringApp^25^ (version 2.2.0) at various confidence levels (0.4 and 0.8), however, 0.8 confidence score-based analysis was the one elected for visualization. PPI network nodes were sized and color-coded based on log fold-change direction and magnitude. Furthermore, network topology metrics were calculated by CytoHubba ^26^, to determine key nodes and candidate hub genes. The Markov Clustering Algorithm was used to build functional clusters out of the differentially expressed genes. To compare gene expression between CF-derived and whole myocardial tissue analyses, we utilised a bulk myocardial RNA-sequencing dataset recently reported by our group ^27^, comprising patients with HOCM (n = 6) and non-failing controls (n = 4). Gene expression patterns were qualitatively evaluated using TPM-normalised expression values and differential expression statistics with FDR ≤ 0.05 as a threshold.

To further examine the cellular specificity of the identified CF signatures, publicly available human HCM single-nucleus RNA-sequencing data were interrogated through the corresponding online interactive portal to the study by Chaffin and colleagues ^28^. Expression of genes was examined across annotated cardiac cell populations, with particular emphasis on fibroblast-enriched clusters.

### Western blotting

As we previously described ^29,30^, Cells and tissues were lysed with a cold RIPA buffer with protease inhibitor (Roche). Lysates were centrifuged for 20 min at 25,000 g at 4°C. The supernatant was quantified using BCA™ Protein Assay Kit (Thermo Scientific). 25 μg of protein lysate per sample was run in 10% gel (Biorad) running system (Life Technologies) in the presence of 1xNuPage MOPS SDS running buffer (Thermo Scientific). Separated proteins were transferred from the SDS polyacrylamide gel onto Whatman® Protran® Nitrocellulose Transfer Membrane (0.2 μm) (Sigma) using a Biorad transfer module, protein transfer buffer (1X NuPage transfer buffer, 10% methanol in dH_2_O). The membrane was incubated for 30 min at RT in 5% skimmed milk blocking solution (1X PBS Tween-20 wash buffer). The blot was then incubated for 2 hrs at RT with primary antibody (**Supplementary Methods**). Blots were washed three times with washing buffer for 15 min, incubated for 1 hr at RT with HRP-labelled secondary antibody (Life Technologies). All incubations and washes were performed using oscillating shakers. Detection was carried out using ECL detection reagents (Life Technologies) and signal was visualized using Amersham Imager 600.

### Histopathology and Immunohistochemistry

As previously described ^29–31^, fixed tissue was processed using an autoprocessor (Leica). 5 μm sections of FFPE tissue sections were deparaffinized in xylene (Sigma, 534056) for 10 mins, hydrated in decreasing concentrations of alcohol (Merck, #100983) and immersed in tap water. 5 μm FFPE sections were deparaffinized in xylene (Sigma, 534056) for 10 mins, hydrated in decreasing concentrations of alcohol (Merck, 100983), and immersed in tap water. Antigen retrieval was performed using 1 mM EDTA (Sigma, 60-00-4) buffer (pH 8) under high pressure and all other incubations were performed at RT using a humidity chamber. Sections were blocked with pre-diluted 2.5 % goat serum (GibcoTM, 16210-064) for 20 min then incubated with primary antibody for 2 hrs. All antibodies were diluted using blocking solution (**Supplementary Methods**). Sections were incubated for 30 min with goat polyclonal anti-rabbit/anti mouse (HRP labelled) secondary antibody and washed thrice prior to staining with DAB + Chromogen for 2 min. Sections were counterstained with hematoxylin (Dako, #S3309), dehydrated through increasing concentrations of ethanol (Merck, #100983) then xylene (Sigma, #534056) before mounting with coverslips using DPX mounting medium (Sigma, 06522).

### Immunocytochemistry

CFs were fixed with 4% paraformaldehyde (Serva, #31628.01) for 20 min at RT and then incubated with chilled absolute methanol for 1 min. Cells were then washed with PBS, incubated with a permeabilization buffer (PBS + 0.5% Triton-100) for 10 min at 4°C, and rinsed with a quenching buffer (100 mM Glycine (Sigma, G8898) in PBS) thrice, 10 min each. Cells were incubated with blocking buffer (PBS + 10% FBS, 7.7 mM NaN3 (Sigma, S2002), 0.1% BSA (Sigma, A2153), 0.2% Triton x-100 (Sigma, T8787), and 0.05% Tween-20 (Sigma, #P1379)) with rocking at 120 rpm for 1 h at RT, and then treated with the antibody diluted in blocking buffer (**Supplementary Methods**). Cells were washed three times,10 min each, with blocking buffer with rocking, and then incubated with the secondary antibody at 1:1000 and kept in the dark for 1 h at RT. Cells were washed with PBS three times with rocking and a mounting medium containing DAPI was added. Images were acquired with confocal microscopy (Zeiss LSM880).

### Cytokine Profiling

Human cytokine arrays (ARY005; RCD Systems) were utilized to investigate the expression of 36 cytokines in the secretome of HOCM-CFs and controls. Array blots were incubated in a blocking buffer for 1 h at RT. 1.5 mL of the sample/antibody mixture was added per chamber, followed by incubation overnight at 2–8°C on a rocking shaker. The washing steps to each blot were performed thrice with 1X wash Buffer at RT. Secondary antibody streptavidin-HRP in array buffer was added at 1:2000 dilution, and blots were incubated for 30 min at RT on a rocking shaker. Blots were washed, followed by the addition of Chemi Reagent Mix for 1 min. Blots were visualized using ECLs and blots were exposed for 1-10 minutes.

### RNA extraction, cDNA synthesis and Real time PCR

RNA was extracted using RNeasy Mini Kit (Qiagen) with an on-column DNase treatment using DNase I Set (RNase-free) kit (zymo research). cDNA was produced using 500 ng RNA with SuperScript IV VILO (Thermo Scientific). PCR reactions were carried out using a QuantStudio™ 5 Real-Time PCR System (Thermo Scientific) under standard conditions. A total of 10μl of reaction mixture contained 1 μl of primer mixture (7.2 μM of both forward and reverse primers), 5 μl of 2x SYBR green Master Mix (Applied biosystems), 2.5 μl of diluted cDNA and dH_2_O to the final reaction volume. Primers sequences are listed in **Supplementary Methods**. Relative mRNA expression was calculated using the 2^−ΔΔCt^ method.

### Statistical analysis

Statistical comparisons between HOCM and control groups were performed according to data type and distribution. Continuous variables were assessed for normality prior to group comparison. Normally distributed data were summarized as mean ± SD and compared using unpaired two-tailed Student’s t-tests. Non-normally distributed data were summarized as median with interquartile range and compared using Mann–Whitney U tests. Categorical clinical variables were summarized as counts and percentages. Densitometric quantification of immunoblots was normalized to GAPDH or β-actin, as appropriate, before comparison between groups.

All statistical analyses were performed using R and GraphPad Prism. Statistical significance was defined as p < 0.05 for targeted validation experiments and q-value/FDR-adjusted p-value ≤ 0.05 for transcriptome-wide and enrichment analyses.

## Results

### 1. Cohort characteristics

The study cohort comprised 12 HOCM patients and 3 controls. The HOCM patients represented a severely symptomatic population (NYHA Class 3/4) with evidence of obstructive septal hypertrophy. All HOCM subjects exhibited significant left ventricular outflow tract (LVOT) obstruction requiring surgical myectomy. Histological examination of myocardial specimens collected from patients were used to further confirm HOCM histological phenotype [myocardial fibrosis and disarray] (**Supplementary Figure 3**).

Clinical phenotyping is summarized in **Table 1**. The cohort showed genetic heterogeneity, with rare heterozygous variants detected in 7 of 12 patients across sarcomeric and non-sarcomeric cardiac genes, including *MYH7, MYBPC3, JPH2, ACTN2, PRKAG2, PTPN11,* and *MYL2*. Because all detected variants were classified as ‘variants of unknown significance’, genotype-specific interpretation of the CF transcriptomic signature was not performed (**Supplementary Table 1)**. Written informed consent was obtained from all patients with HCM before inclusion in the study. The study was approved by the Aswan Heart Centre Ethical Committee ([20130405MYFAHC_CMR_20130330]). Control CFs were obtained from the Magdi Yacoub Institute, UK, through the heart donation programme under a MTA, with ethical approval from the Royal Brompton Hospital/Brompton and Harefield Trust Ethics Committee (Research Ethics Committee approval 10/H0724/18).

**Table 1:** Clinical and demographic characteristics of the study cohort.

| Clinical Parameters | HOCM (n = 12) | Controls (n=3) |
| --- | --- | --- |
| Gender, Male/Female | 6 Males/6 Females | 3 males |
| Age (SD) | n = 12<br>47 ± 19.2 | n=3<br>60 ± 4.9 |
| Body surface area BSA. m<sup>2</sup> | n = 12<br>1.8076 ± 0.37 | NA |
| Body mass index | n = 12<br>30.3 ± 8.3 | NA |
| NYHA class | Class 2 (n = 3) (25%)<br>Class 3 (n = 8) (66.7%)<br>Class 4 (n = 1) (8.3%) | NA |
| Max Septal wall thickness by Echocardiography (cm) | n = 12<br>2.43 ± 0.437 | NA |
| LVOTO resting gradient by Echocardiography (mmHg) | n = 12<br>103.92 ± 40.76 | NA |
| Left ventricular mass by CMR (gram) | n = 10<br>173.5 (165.75–220) | NA |
| Left ventricular end systolic volume indexed by CMR (ml) | n = 12<br>18 ± 8 | NA |
| Left ventricular end diastolic volume indexed by CMR (ml) | n = 12<br>81 ± 12 | NA |
| Left ventricular Ejection Fraction% | n = 12<br>78.3 ± 7.5 | NA |
| BNP (pg/mL) | n=11<br>1711 (304-6176) | NA |
| Cardiac Troponin I (ng/mL) | n = 11<br>0.011 (0.008–0.014) | NA |

NA: not available to control CFs

### 2. Transcriptomic Profiling Reveals Inflammatory Activation and Selective ECM Remodelling in HOCM-CFs

CFs isolates were assessed using marker-based approaches. Immunocytochemistry and flow cytometry confirmed enrichment of fibroblast-associated markers, including vimentin (VIM), CD90 and α-SMA, with low expression of endothelial, immune and myocyte-associated markers (**Supplementary Figure 2**). RNA-seq data were used as an additional quality-control step to assess lineage-marker expression across the cultured isolates. Transcriptomic assessment of lineage markers showed high expression of fibroblast-associated genes, including *VIM, COL1A1, α-SMA/ACTA2, periostin (POSTN),* Platelet-Derived Growth Factor Receptor Alpha (*PDGFRA)*, Transcription Factor 21 *(TCF21)* and CD90 (*THY1)*, with minimal expression of non-fibroblast lineage markers *PECAM1*/CD31, *PTPRC*/CD45 and Desmin (*DES)* (**Supplementary Figure 3**).

Comparative transcriptomic analysis of 12 HOCM-CFs and 3 control counterparts revealed 265 significant DEGs (**Figure 1A, Supplementary Data 1).** Unsupervised clustering and Principal Component Analysis (PCA) showed a segregation between HOCM-CFs and control samples, supporting the relevance of the filtered DEG signature for downstream enrichment and candidate-gene analyses (**Supplementary Figure 4A to 4D**). Expression-density analysis further indicated a difference in the DEG expression distribution between HOCM-CFs and controls (**Supplementary Figure 4E)**. The top 20 significantly enriched Reactome pathways (according to Benjamini-Hochberg adjusted p-value) clustered into three functional categories: immune system (7 pathways), structure and structural components (6 pathways), and signaling (7 pathways) (**Figure 1B, Supplementary Data 2**). Of note, the top three significantly enriched pathways were ‘*Interleukin-4 and Interleukin-13 Signaling’*, ‘*Extracellular Matrix Organization’*, and ‘*Chemokine Receptor Bind Chemokines’*, underscoring a prominent immunoinflammatory and structural remodeling signature in HOCM-CFs (**Figure 1B**).

**Figure 1:**
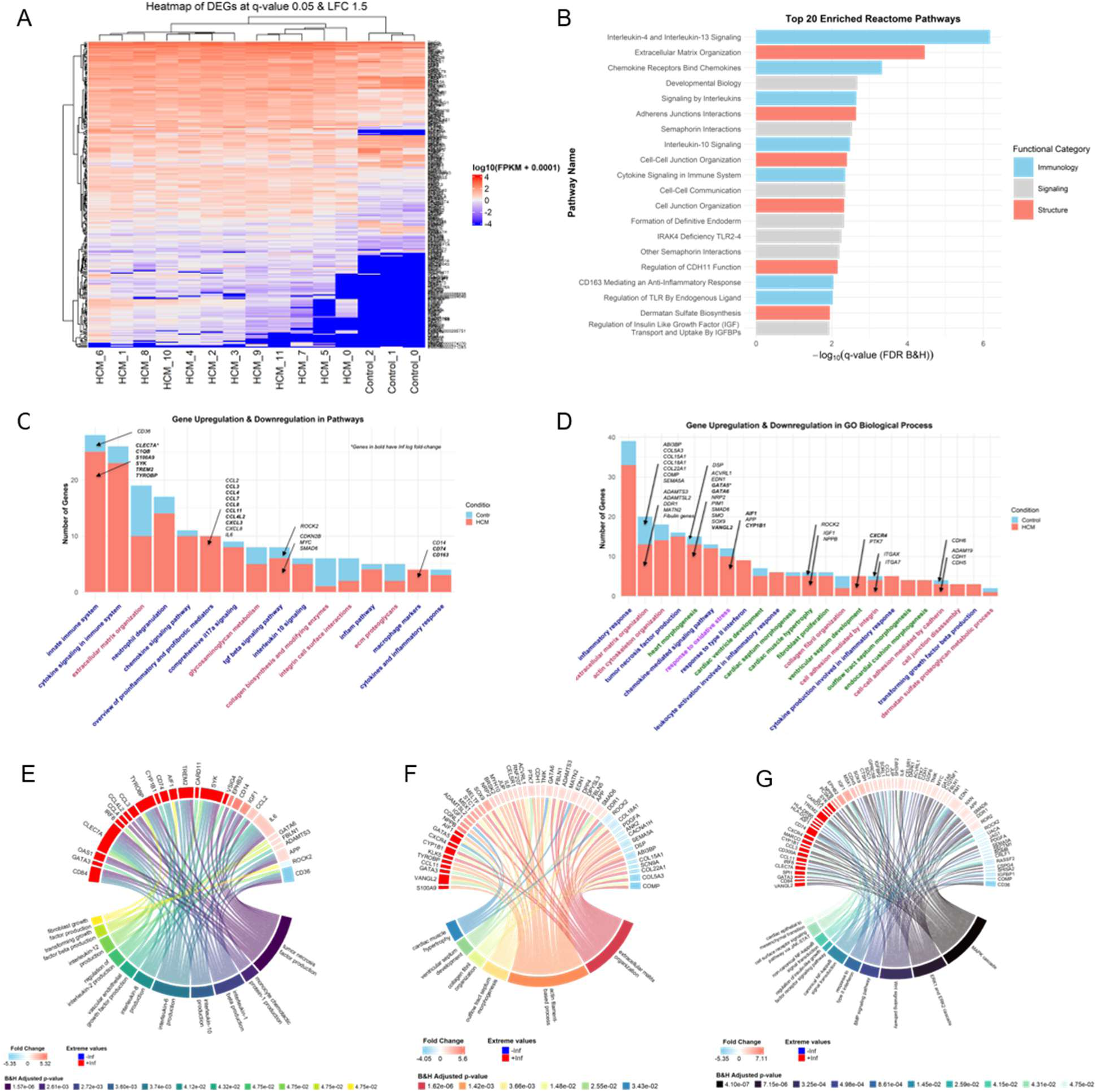
Transcriptome signature of HOCM-CFs and associated pathways. **A:** Heatmap of log-transformed FPKM values for DEGs across control and HOCM samples. Rows represent genes and columns represent biological replicates. **B:** Bar plot of the top 20 Reactome enriched pathways, order according to the Benjamini-Hochberg adjusted p-value. **C-D:** Stacked bar plot of select enriched pathways of interest in C and GO biological processes of interest in D, showing the proportion of upregulated and downregulated genes in each term. Immunology-related terms are in blue, structure-related terms are in magenta, cardiac terms are in green and oxidative stress-related terms are in purple. **E-G:** Circos plot of select enriched interleukin and growth factor production pathways in **E**, structure-related pathways in **F**, and signaling pathways in **G**.

Analysis of the directionality of DEGs included selected enriched pathways and GO biological processes showed that the two most represented terms were ‘inflammatory response’ and ‘extracellular matrix organization’, indicating that both inflammatory signalling and ECM remodelling are central features of the HOCM-CFs transcriptomic signature (**Figures 1C–D, Supplementary Data 3-4**). The ‘*inflammatory response’* term was mainly composed of upregulated genes, with a relatively smaller contribution from downregulated genes, consistent with enhanced inflammatory activation in HOCM-CFs. ‘*Extracellular matrix organization’* contained upregulated and downregulated genes, suggesting active ECM remodelling rather than uniform ECM induction or suppression. Across the remaining enriched terms, immune- and stress-related processes, including leukocyte activation, cytokine/chemokine signalling, oxidative stress responses and inflammatory mediator production, generally showed a tendency toward upregulation, whereas structural, adhesion, morphogenesis and matrix-associated processes showed more heterogeneous directionality.

Circos plots were used to visualise the contribution of individual DEGs to selected enriched pathway clusters and gene overlap between biological processes. In contrast to the broader pathway categories shown in **Figure 1C** and **1D**, circos plots focused on more specific pathways within each biological cluster to enable higher-resolution visualisation of pathway-specific gene contributions (**Figures 1E-G, Supplementary Data 5-6**). Within Inflammation-related pathways, representing enriched interleukin and growth factor production pathways, several key upregulated genes were identified including *ILC, CCL2, CCL8, CLEC7A, SYK, TYROBP, CD36*, and *CD84* (**Figure 1E**). Within ECM organization-related pathways, upregulated genes included *ADAM9, ADAM33, ADAMTS3, MATN2, PTK7, SEMA5A, FBLN1, FBLN5* and *ABI3BP*, while *COL5A3, COL22A1, COL18A1*, and *COMP* were downregulated (**Figure 1F**). Within growth factor signaling and fibroblasts activation-related pathways, upregulated genes included *EDN1, TREM2, IGF1, CD74, ROR2,* and *APP* (**Figure 1G).**

### 3. Network Analysis Reveals Inflammatory Hub Genes Within the HOCM-CFs DEG Signature

To investigate the potential functional connections within the DEGs, protein-protein interaction (PPI) network analysis of all 265 DEGs revealed multiple interactions between ECM- and inflammation-related proteins within the HOCM-CFs transcriptome signature (**Figure 2A, Supplementary Data 7-9**). Markov Clustering Algorithm clustering (inflation parameter = 5) identified functionally coherent protein modules, predominantly composed of upregulated genes. Networks were visualised using a high-confidence interaction threshold (STRING confidence score = *0.8*), which preferentially retained robust interactions; ECM/collagen-associated subnetworks were detectable only at lower confidence thresholds (default confidence score = *0.4*) (**Supplementary Figure 5**). Network topology analysis consistently ranked four genes among the top 15 nodes across the Maximal Clique Centrality (MCC), degree, and betweenness centrality metrics: *CXCR4*, *IL6*, *CD74*, and *TYROBP*. The *CXCR4* node had the highest degree within the chemokine activity module. The cross-talk between the two main immune response modules of the network was through the CXCR4/CD74 axis. These proteins have the highest overall betweenness, suggesting their potential role as critical control points. Moreover, *CXCR4* had the highest degree within the chemokine activity module. *IL6* node occupied a central position within the cytokine/chemokine signaling module, as it is one of the key pro-inflammatory cytokines known to be markedly elevated in myocardial pathological conditions associated with fibrosis^32^. *TYROBP* connected multiple immune-inflammatory modules, consistent with its role in myeloid-cell activation and upstream regulation of inflammatory signaling (**Figure 2B, Table 3 and Supplementary Data 5 and 6**).

**Figure 2.**
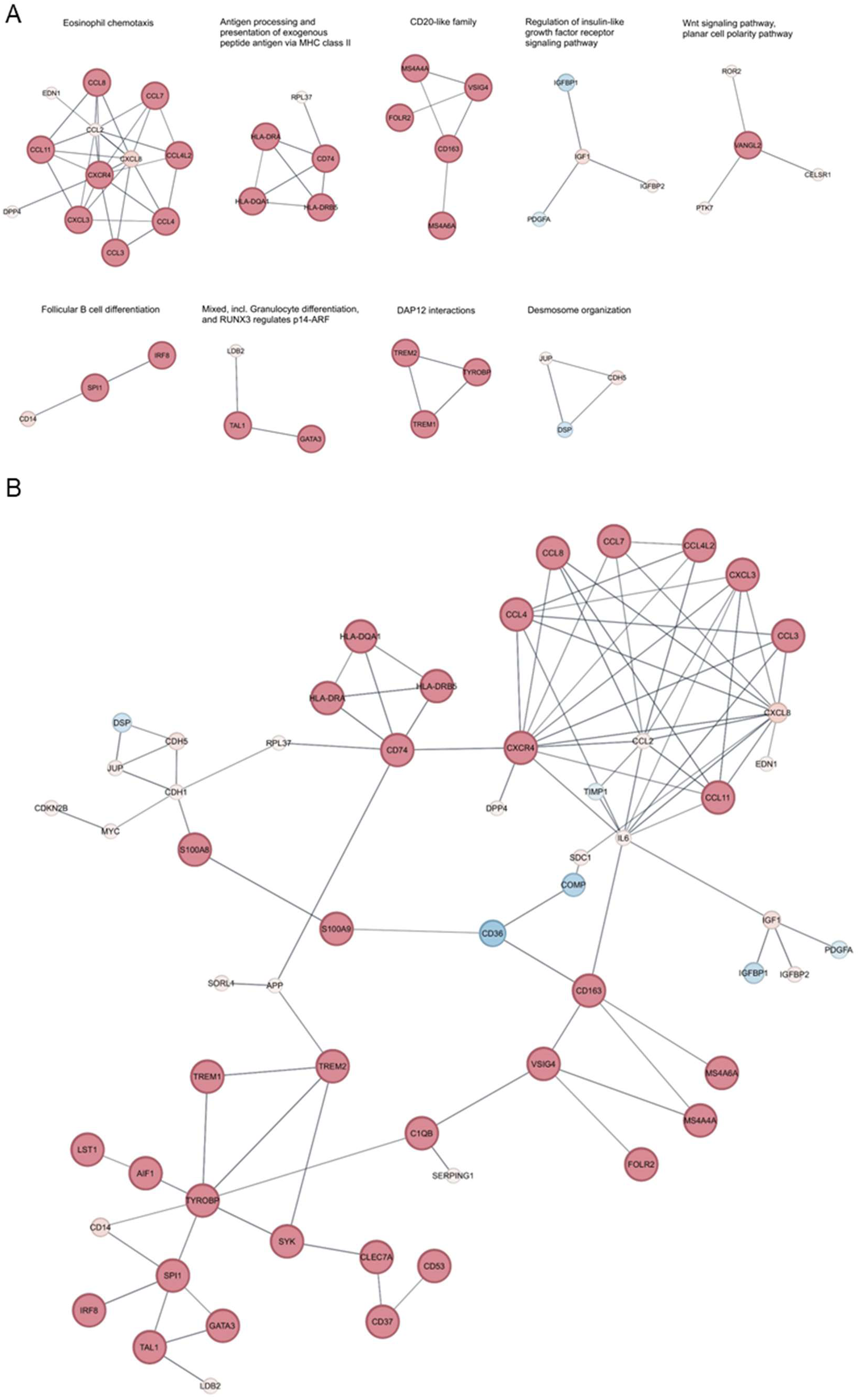
Protein interactome network in HOCM-CFs signature. **A:** Network clusters formed by the DEGs of the Tuxedo protocol using Cytoscape StringApp. **B:** Protein-protein interaction (PPI) network of the DEGs identified by the Tuxedo protocol. The nodes are sized <u>based on fold-change magnitude,</u> and colored based on fold change <u>direction</u>. Upregulated genes are red and downregulated genes are blue.

### 4. CFs-focused profiling resolves inflammatory and ECM-remodelling signatures not captured by whole tissue RNA-seq

Based on the DEG distribution and GO enrichment profile, we focused the genes encompassing the two most represented biological processes: ‘*inflammatory response’* and ‘*extracellular matrix organization’*. This analysis was intended to define the gene-level contributors to these dominant GO terms and to assess whether the CFs-associated inflammatory and ECM-remodelling signatures were qualitatively reflected in independent HOCM myocardial tissue RNA-seq data. Within the ‘*inflammatory response’* GO term, HOCM-CFs showed a predominantly upregulated pattern compared with control CFs (**Figure 3A**). This gene set included inflammatory, chemokine, immune-associated, and signaling mediators such as *IL6, CCL2, CCL4, CCL11, CXCR4, TYROBP, TREM2, CD14, CD36, CD300A, CLEC7A, SYK, APP, IGF1, FOLR2, LYZ*, and *CXCL8*, supporting the enrichment of cytokine/chemokine signaling, immune-associated receptor activity, and inflammatory mediator production.

**Figure 3:**
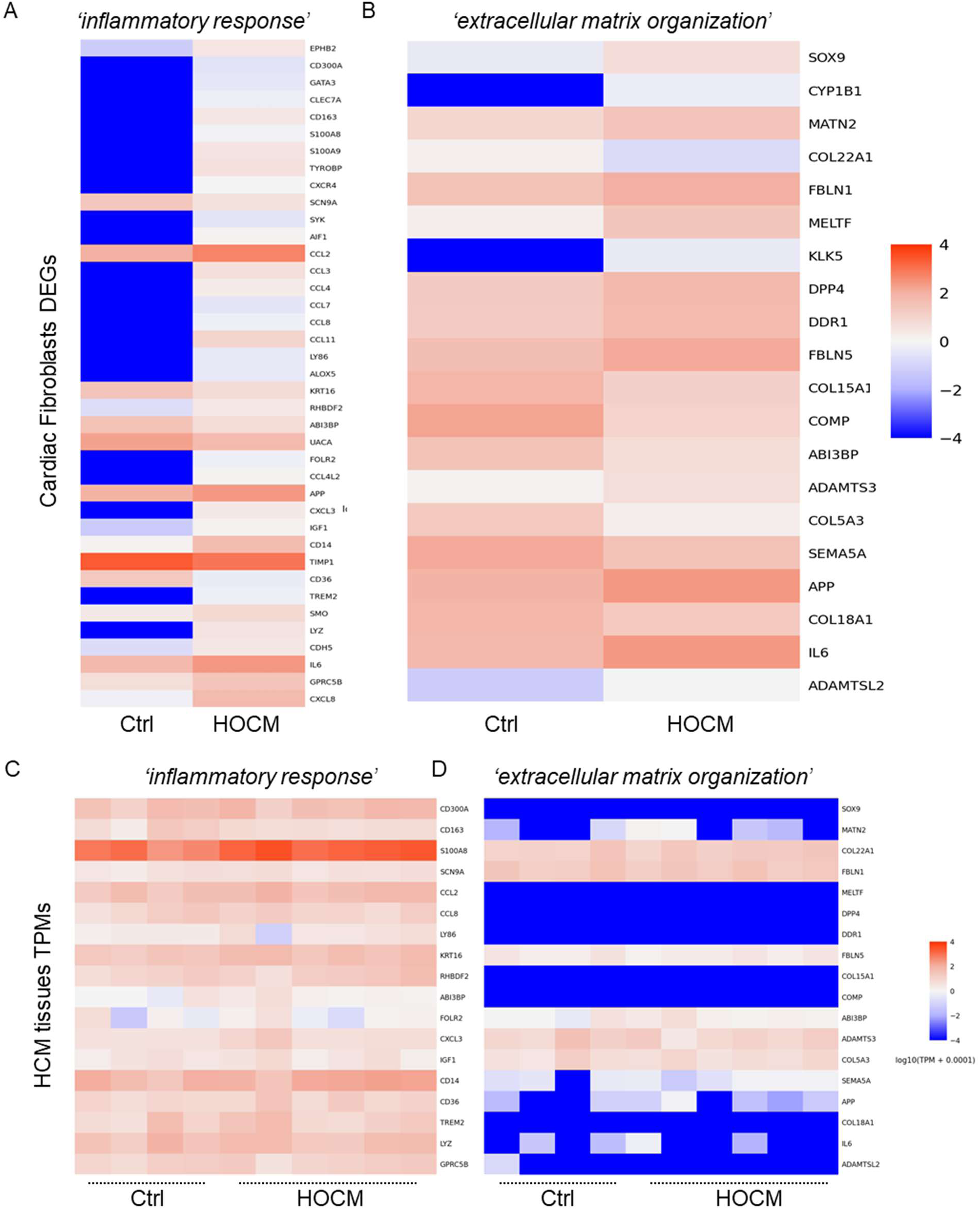
Comparison of inflammatory signaling and ECM-remodeling gene signatures in HOCM-CFs and HOCM myocardial tissue. Heatmaps showing expression patterns of genes assigned to the inflammatory response/signaling and extracellular matrix organization/remodeling GO terms in cardiac fibroblasts and myocardial tissue datasets. **A and B:** The mean expression of control versus HOCM-CFs. **C and D:** The expression across individual control and HOCM myocardial tissue samples. Genes were selected from enriched GO terms identified in the HOCM-CFs DEG analysis. The inflammatory gene set includes cytokines, chemokines, immune-associated receptors, and inflammatory signaling mediators, while the ECM-remodeling gene set includes collagens, fibulins, proteases, matrix-associated glycoproteins, and cell–matrix interaction genes. Expression values are shown as log10(TPM + 0.0001).

Genes assigned to the ‘*extracellular matrix organization’* GO term showed a more heterogeneous and bidirectional pattern in HOCM-CFs, consistent with selective ECM remodelling rather than uniform matrix induction (**Figure 3B**). Upregulated ECM-remodelling and matricellular genes included *MATN2, FBLN1, FBLN5, DDR1, ADAMTS3, APP, IL6*, and *ADAMTSL2*, whereas several structural or matrix-associated components, including *COL5A3, COL15A1, COL18A1, COL22A1, COMP, ABI3BP*, and *SOX9*, showed a downregulation pattern in HOCM-CFs. These findings support the interpretation that HOCM-CFs undergo ECM reorganization involving elastic fibre-associated glycoproteins, matrix-interacting receptors, protease-related remodelling, and altered structural matrix components.

To assess tissue-level overlap, the 265 CFs-HOCM DEGs were matched to previously published RNA-seq data from HOCM myocardial tissue and control samples ^27^. Of these, 254 genes were detected in the tissue TPM dataset. However, statistical comparison of 4 control tissue samples against 6 HOCM tissue samples showed that none of the overlapping genes reached significance after FDR correction (**Figure 3C and 3D**; **Supplementary Data 10**). To further contextualize the HOCM-CF signature at the cellular level, selected DEGs were interrogated in the human cardiomyopathy single-nucleus RNA-sequencing atlas reported by Chaffin and colleagues ^28^. Analysis of myocardial nuclei from 16 non-failing, 15 HCM and 11 DCM hearts demonstrated that several inflammatory and ECM-remodelling genes identified in HOCM-CFs were detectable in diseased myocardium, including genes associated with cytokine signalling, matrix remodelling and fibroblast activation (**Supplementary Figure 6**). Although the atlas analysis was exploratory and performed at the whole-tissue level, owing to limitations of the portal-based interrogation, the findings provide independent support for the biological relevance of the inflammatory and ECM-remodelling programmes identified in HOCM-CFs. This reflects the importance of CFs-focused profiling for uncovering fibroblast-specific disease mechanisms in HOCM.

### 5. Protein validation confirms inflammatory secretion and selective ECM remodelling in HOCM

To determine whether the transcriptomic HOCM-CFs signature translated into quantifiable protein changes, we prioritised inflammatory cytokine/chemokine signalling and ECM/matrisome remodelling.

To validate the inflammatory component of the HOCM-CFs signature, cytokine profiling was performed on conditioned media collected from HOCM-CFs and control-CFs using a cytokines array screening 36 inflammatory cytokines and chemokines. This analysis identified IL6 and CCL2 as significantly increased in HOCM-CFs conditioned media compared with controls, supporting an enhanced inflammatory secretory phenotype in HOCM-derived CFs. Additional inflammatory mediators, including IL8, CXCL1, MIF and PAI-1, showed increased expression trends in HOCM-CFs conditioned media, although these did not reach statistical significance (**Figure 4A**). Interestingly, IL8 has recently been implicated in HCM pathogenesis, with CF-derived IL8 reported to promote fibrosis and tissue stiffening through CXCR1 signaling ^33^, suggesting that this pathway may contribute to the pro-inflammatory and pro-fibrotic phenotype observed in HOCM-CFs.

These findings were confirmed at the cellular level by immunocytochemistry, which showed increased IL6 and CCL2-associated signal in HOCM-CFs compared with control-CFs (**Figure 4B**). qPCR and Immunoblotting analyses of CF lysates confirmed significantly higher IL6 and CCL2 expression in HOCM-CFs (**Figure 4C** and **4D**). Consistent with the CF-derived inflammatory signature, immunoblotting of HOCM myocardial tissue lysates showed increased IL6 and CCL2 protein expression (**Figure 4E**). Immunohistochemistry of IL6 in HOCM tissues showed an evident increased expression compared to control tissues, though the expression pattern did not appear to be specific to stromal cells (**Figure 4F**). CCL2 tissue localization was not assessed by immunohistochemistry because of technical limitations with staining optimization.

**Figure 4:**
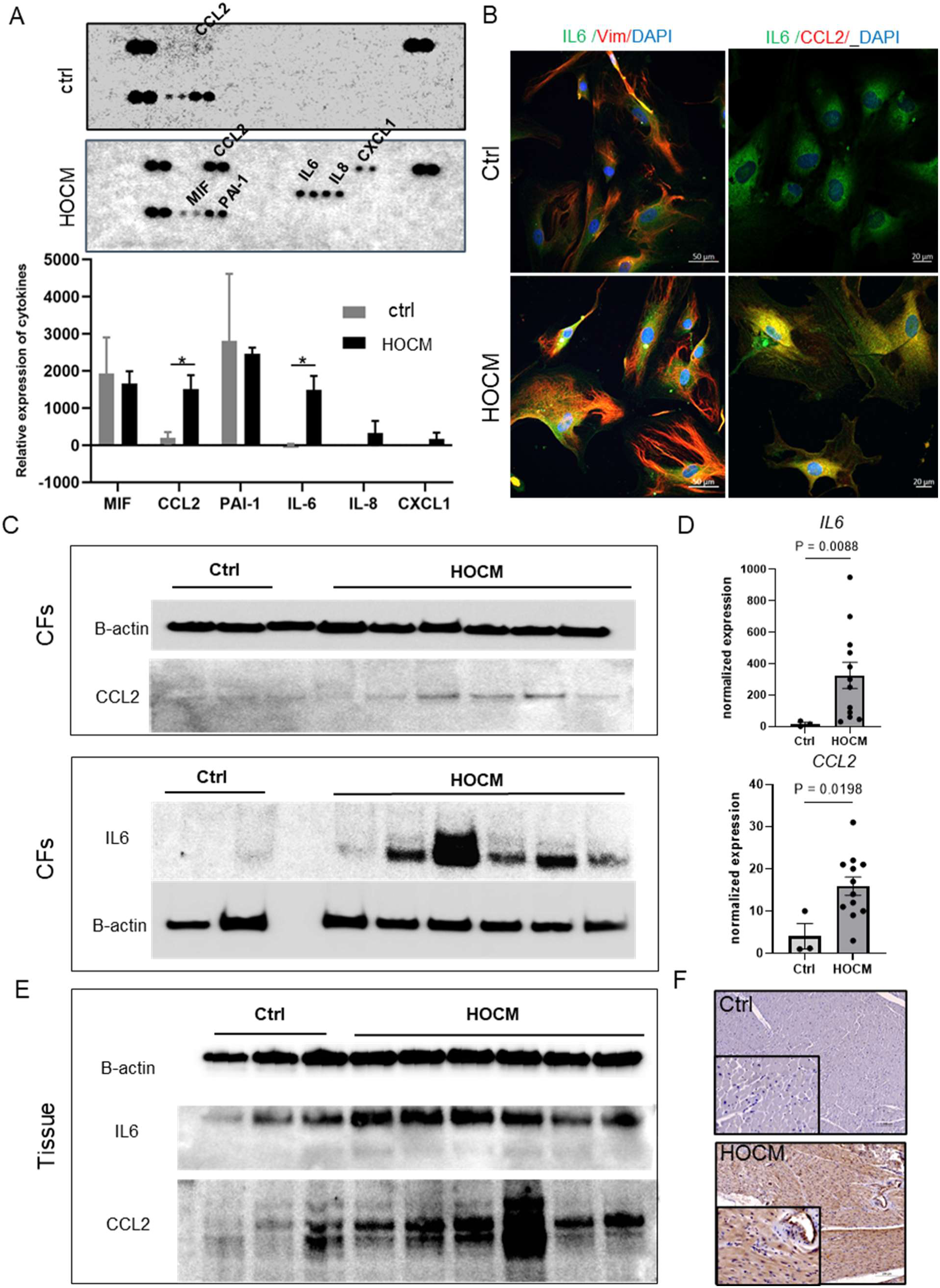
Validation of inflammation-related genes in HOCM CFs and myocardial tissues. **A:** Cytokines and chemokines profiling for the conditioned media collected from HOCM-CFs (n=12) vs Ctrls (n=3), showing the significant upregulation of *ILC* and *CCL2* in HOCM-CFs. B: Immunocytochemical staining of *ILC* and *CCL2* in HOCM CFs (n=6) vs control counterparts (n=3). Scale bars are 50µm and 20µm. **C**: Immunoblots for *CCL2* and *ILC* (normalized to B-actin) in HOCM-CFs lysates (n=6) vs controls (n=3). **D**: qPCR analysis for mRNA of *CCL2* and *ILC* (normalized to B-actin), in HOCM-CFs (n=12) vs controls (n=3). **E**: Immunoblots for CCL2 and IL6 (normalized to B-actin) in HOCM tissue lysates (n=6) vs controls (n=3). **F**: Immunohistochemistry staining for *ILC* in HOCM tissues (n=12) vs ctrls (n=3). Scale bars are 100µm.

To validate the ECM/matrisome component of the HOCM-CFs signature, representative ECM-related DEGs with expression direction were selected for targeted assessment, including *FBLN1* (up), *CSPG4* (down), *ABI3BP* (down) and *COL5A3* (down). qPCR analysis confirmed the expression pattern and differential expression of these genes in HOCM-CFs compared with controls (**Figure 5A**). At the protein level, immunoblotting of CFs lysates showed increased expression of selected ECM-remodelling proteins, including FBLN1, FBLN5, and MMP9, in HOCM-CFs, whereas COLV did not show a clear difference between HOCM- and control-derived CFs (**Figure 5B–C**). Although MMP9 was not included in the DEGs, it was included in the validation panel because of its known matrix-remodelling activity and its potential role in COLV turnover ^34^.

**Figure 5:**
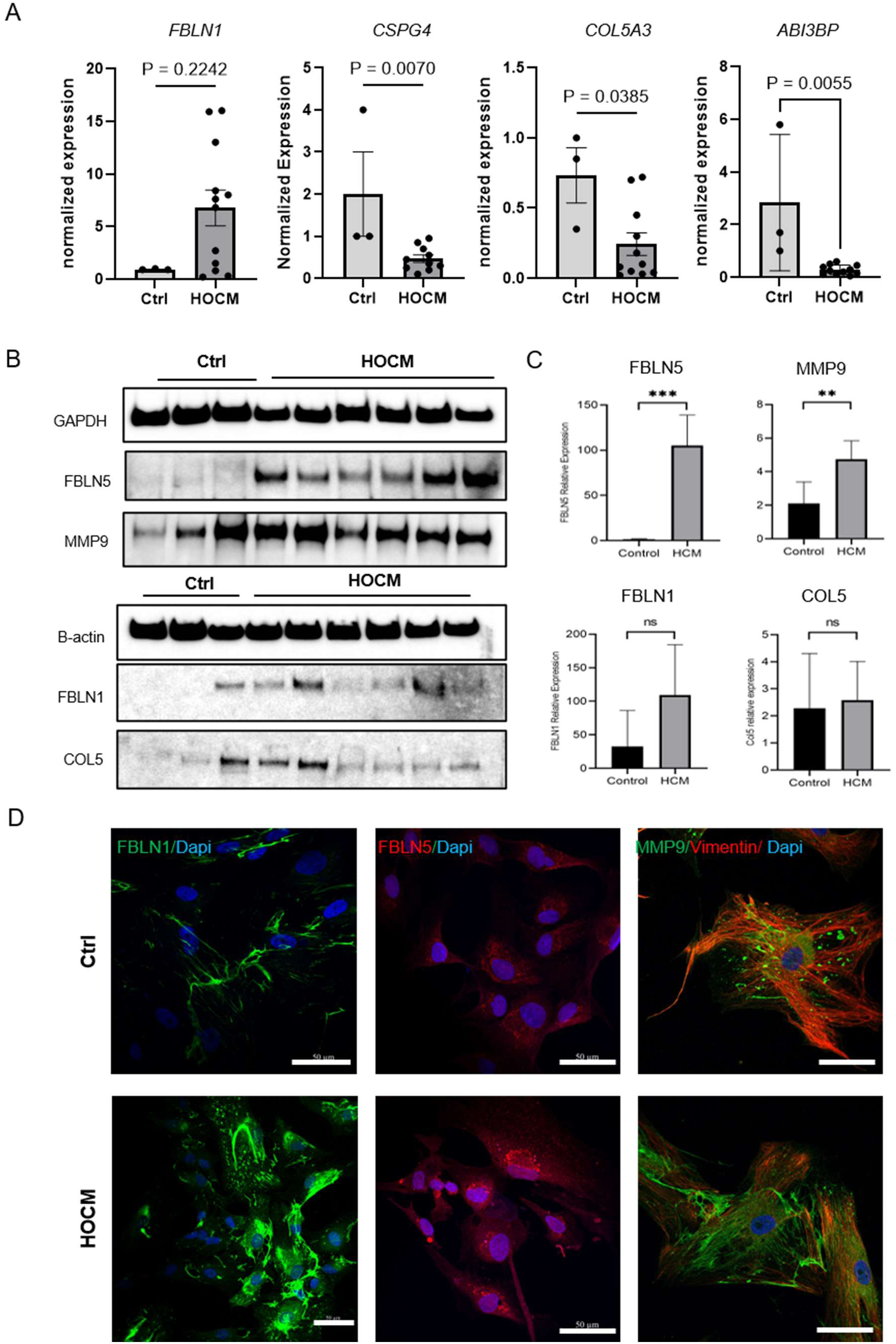
Validation of ECM-related genes in HOCM CFs. A: qPCR analysis for mRNA of *FBLN1*, *CSPG4*, *ABI3BP* and *COL5A3* (normalized to B-actin), in HOCM-CFs (n=12) vs controls (n=3). **B:** Immunoblots for MMP9, FBLN1, FBLN5 and ColV in HOCM CFs lysates (n=6) vs controls (n=3). **C**: Quantification of ECM markers in CFs-immunoblots normalized to GAPDH or B-actin. **D:** Immunocytochemistry of MMP9 and vimentin, FBLN1 and FBLN5 in HOCM-CFs (n=6) vs ctrls (n=3).

Immunocytochemistry supported the altered ECM-remodelling profile of HOCM-CFs, showing stronger cellular/pericellular staining for FBLN1, FBLN5, and MMP9 compared with control CFs (**Figure 5D**). These findings suggest that HOCM-CFs acquire a matrix-remodelling phenotype characterised by increased elastic fibre-associated glycoproteins and matrix-degrading activity, rather than a uniform increase in all collagen-associated ECM components.

Consistent with the CFs-level findings, immunoblotting of myocardial tissue lysates confirmed increased FBLN1, FBLN5 and MMP9 expression in HOCM tissue compared with controls, while COLV did not show a significant increase by immunoblotting (**Figure 6A and 6B**). Immunohistochemical analysis showed increased MMP9 and FBLN1 staining, together with a reduced COLV staining, in HOCM myocardium compared with controls (**Figure 6C**). Notably, FBLN1 and COLV staining appeared predominantly stromal in HOCM tissue, whereas MMP9 displayed both stromal and cardiomyocyte-associated localisation. These validation experiments support the transcriptomic prediction that HOCM-CFs are characterised by a combined inflammatory and ECM-remodelling phenotype, with disease-associated changes detectable in cultured patient-derived CFs and in HOCM myocardial tissue.

**Figure 6:**
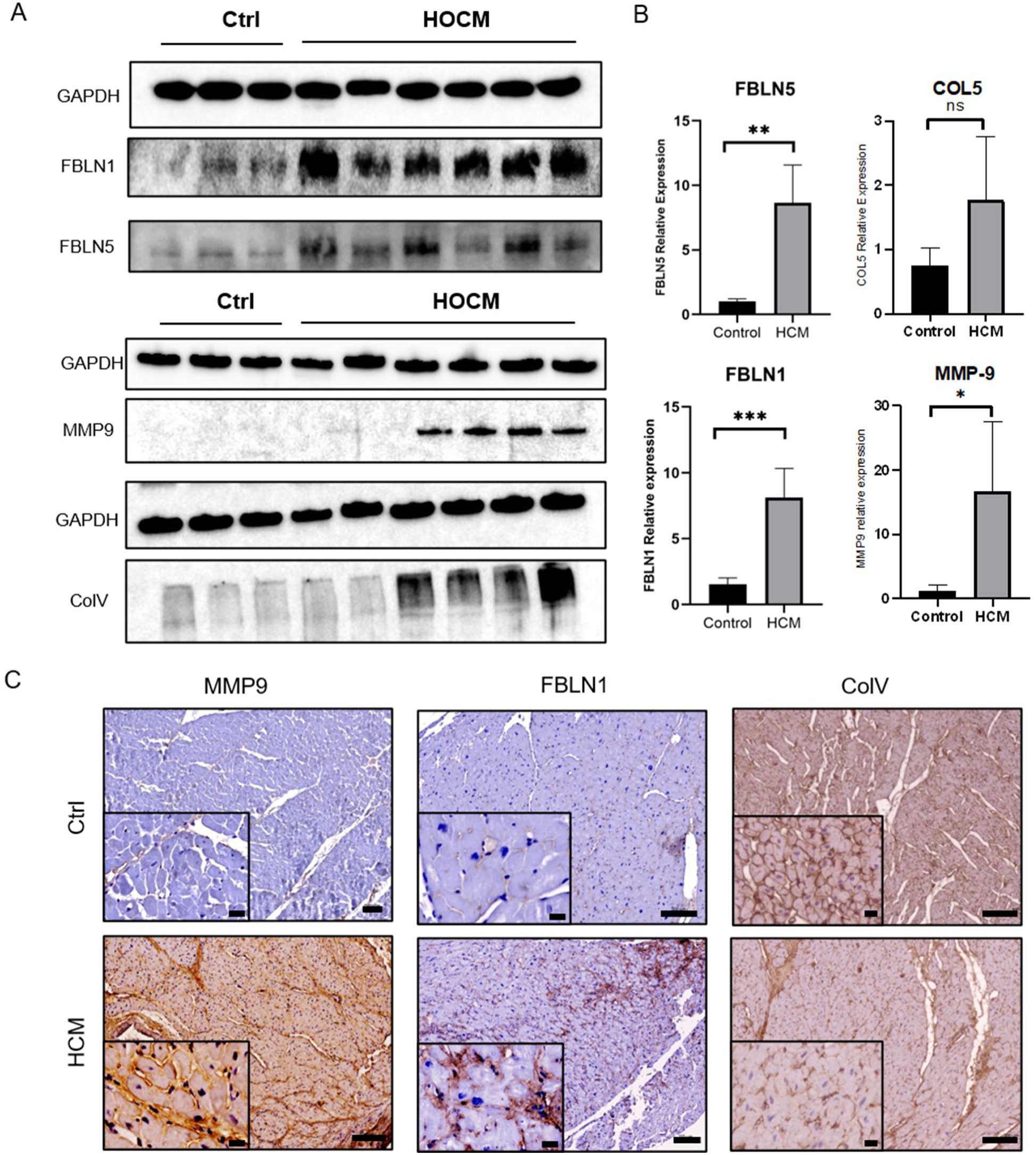
Validation of ECM-related genes in HOCM myocardial tissue. **A:** Immunoblots for *MMPS, FBLN1, FBLN5* and *COLV* in tissue lysates of HOCM myocardium (n=6) vs controls (n=3). B: Quantification of ECM markers in tissue-immunoblots normalized to GAPDH. **C:** Immunohistochemistry staining for *FBLN1, MMPS* and *COLV* in HOCM tissues (n=12) vs ctrls (n=3). Scale bars are 50µm.

As an exploratory assessment of whether the inflammatory and ECM-remodelling mediators identified in HOCM-CFs remain responsive to profibrotic stimulation, cultured HOCM- and control CFs were treated with TGFβ1. Cytokine/chemokine profiling of conditioned media from TGFβ1-treated HOCM-CFs showed increased IL6 and CCL2, while immunoblotting supported increased IL6 protein expression in TGFβ1-treated control CFs (**Supplementary Figure 7A–B**). In parallel, immunoblotting of matrix-associated proteins showed altered expression patterns of COLV, FBLN5, and MMP9 following TGFβ1 exposure (**Supplementary Figure 7C**). These findings suggest that inflammatory and ECM-remodelling mediators remain inducible under profibrotic stimulation.

## Discussion

This study presents a disease-specific transcriptomic state in HOCM patients-derived CFs, characterized by ECM structural remodeling, altered cell–cell and cell–ECM communication, and inflammatory and profibrotic signaling. This signature did not reflect uniform activation of structural ECM genes, but rather a remodeling program involving fibulins, proteases, integrins, matricellular proteins, and a subset of downregulated collagens.

Although interstitial fibrosis has been known as a hallmark of HCM ^35^, the underlying mechanisms and the associated ECM remodeling have not been adequately investigated. We identified groups of CFs-associated ECM gene families with distinct patterns of expression that potentially contribute to cardiac fibrosis and changes in ECM composition. Although most CFs-derived DEGs were detectable in the bulk myocardial tissue RNA-seq dataset, none reached statistical significance after FDR correction. This suggests that the CFs-derived inflammatory and ECM-remodeling program is not statistically resolved at the individual gene level in bulk tissue, likely due to dilution by myocardial cellular heterogeneity. In contrast, profiling of patient-derived CFs provided cell-type resolution to uncover these disease-associated fibroblast programs.

### ECM remodeling extends beyond collagen upregulation

The collagen genes identified within the HOCM-CFs DEG signature were predominantly downregulated, including fibril-forming collagens (COL5A3), multiplexins (COL15A1, COL18A1), and fibril-associated collagens (COL21A1, and COL22A1), most of which have been studied during tissue fibrosis in non-cardiac systems ^36–39^. Collagens I and III did not appear in the CFs-associated signature, despite their reported relevance to myocardial interstitial fibrosis ^40^. Nonetheless, fibrosis reflects the net outcome of matrix synthesis, degradation and crosslinking, rather than transcriptional upregulation of all collagen genes at a single time point. The upregulation of fibulins, proteases, integrins, and matricellular genes suggests that HOCM-CFs may contribute to matrix reorganization and alter ECM quality and turnover in HOCM.

One of the proteoglycans family, *CSPG4*, was downregulated in HOCM-CFs, which is a constituent of the cardiac conduction system ^41^, and has been reported to prevent sympathetic axon regeneration after myocardial infarction ^42^. On the other hand, members of the fibulins glycoproteins family (*FBLN1* and *FBLN5*) and various proteases (such as *ADAM33* and *ADAM19*), were significantly upregulated in HOCM-CFs. While *FBLN1* and *FBLN5* functions have been related to vascular pathologies ^43^, their role in myocardial ECM remodeling has not been studied. We have recently reported that *FBLN2*, which has common binding partners with *FBLN1* and *FBLN5* ^44,45^, is upregulated in the cardiomyocytes and the circulation of HOCM patients, however, protein expression in CFs did not significantly change ^13^; an observation that was further confirmed by our transcriptome signature of HOCM-CFs. In relation to ECM remodeling, the proteases encompassed in CFs signature, such as ADAM19 and ADAM33 have several targets and reported roles in myocardial and vascular pathologies, in particular, shedding of membrane proteins and growth factors. Increased MMP9 expression in HOCM-CFs suggests that post-transcriptional or culture-context-dependent regulation of matrix remodeling enzymes may also contribute to the HOCM-CFs phenotype. Of interest, TIMP1, an inhibitor for MMP activity ^46^, was downregulated in HOCM-CFs, whereas S100A8 and S100A9, which have been linked to inflammatory MMP9 induction in fibroblasts, were upregulated. ^47^, supporting a potential imbalance in ECM turnover in HOCM-CFs.

### Altered cell–matrix communication in HOCM-CFs

Integrins are crucial elements for cell-ECM interaction ^48^. Our data identified the downregulation of *ITGA7* which encodes a laminin-binding integrin alpha subunit when heterodimerized with beta integrin partners, and upregulation of *ITGAX*, which has been linked to fibrinogen and complement-associated binding functions ^49^. This suggests altered CFs–ECM interaction profiles in HOCM-CFs, which may influence fibroblast behavior, adhesion, and response to matrix cues ^48^.

### Inflammatory and profibrotic signaling in HOCM-CFs

Our data identified an inflammatory signature that comprises drivers for inflammation, fibrosis, and immune cells recruitment in HOCM-CFs. Of which, IL6 induces the proliferation of CFs through autocrine and paracrine (from cardiomyocytes and resident immune cells) effects, a mechanism mediated by Angiotensin II and TGFβ1 signaling, that could lead to myocardial damage and fibrosis ^50,51^. Furthermore, deletion of IL6 may attenuate the hypertrophy-related machineries in mouse hearts via a Ca^2+^/calmodulin-dependent protein kinase regulation ^52,53^. Although *WNT5A* has been suggested to induce IL6-associated fibrosis ^54,55^, HOCM-CFs signature identified a significant downregulation of *WNT5B*, with no differential expression of *WNT5A*.

The responsiveness of IL6 and CCL2 to TGFβ1 stimulation supports functional overlap between profibrotic signaling and inflammatory mediator expression in CFs. These data suggest that inflammatory cytokine/chemokine expression remains inducible under profibrotic stimulation and may contribute to the coupling between fibrosis-associated signaling and inflammatory remodeling. We identified *IGF-1* and *PDGFA, which* exhibited relevance to cardiac fibrosis in mouse models ^56,57^. Consistent with this network-level organization, PPI analysis identified *CXCR4*, *IL6*, *CD74*, and *TYROBP* as recurrent hub nodes, suggesting that the inflammatory component of the HOCM-CFs signature is structured around interconnected chemokine, cytokine and myeloid-associated signaling modules. Of interest, a recent study reported a fibroblast-driven MYC–CXCL1–CXCR2 axis in pressure overload-induced cardiac dysfunction and HF development ^58^. Although our HOCM-CFs signature did not identify the same CXCL1–CXCR2 axis, it showed upregulation of related chemokine-associated mediators, including *CXCR4*, *CXCL3*, and *CXCL8*, together with increased *MYC* expression. This suggests that MYC-associated transcriptional programs may contribute to parallel inflammatory and chemokine signaling cascades in HOCM-CFs, potentially reflecting disease-context-specific fibroblast inflammatory remodeling. The dual identification of *CD74* and *CXCR4* as hub nodes highlighted the nature of CFs state to sustain a pro-fibrotic status and CFs survival, as reported to occur via AKT activation ^59^.

These findings shift the interpretation of HOCM-CFs involvement from a purely collagen-producing fibroblast model toward a broader ECM-organization and inflammatory-remodeling fibroblast state. This has potential therapeutic implications, suggesting that fibroblast-directed strategies in HOCM need to consider inflammatory signaling, cell–matrix communication, and ECM turnover alongside classical anti-fibrotic approaches (**Figure 7**).

**Figure 7:**
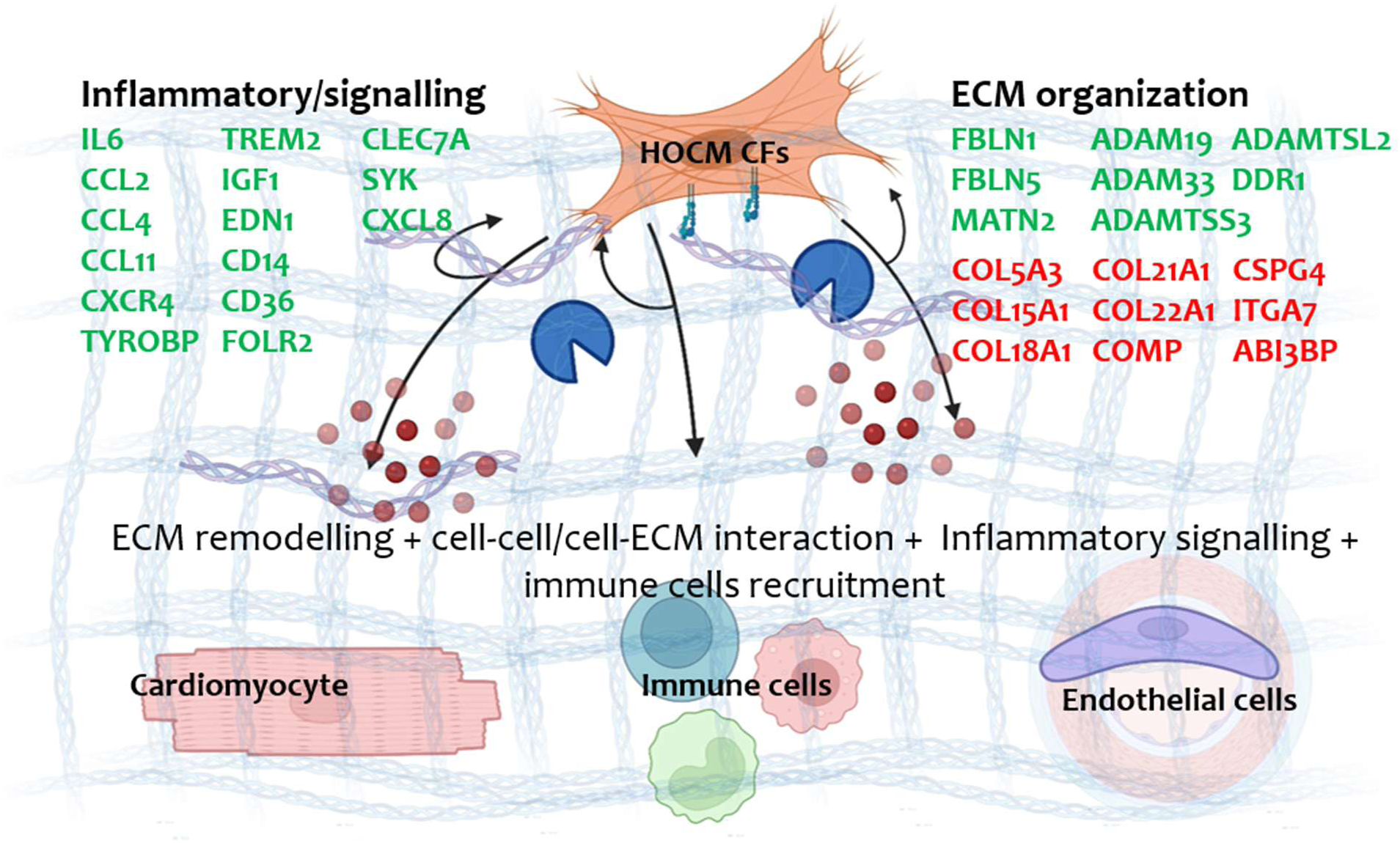
Proposed functional model of the inflammatory-signaling and ECM-remodelling state of HOCM cardiac fibroblasts. Schematic diagram of HOCM-CFs signature and their potential effect on cardiac components, inferred from transcriptomic, pathway, and validation analyses. The model proposes that HOCM-CFs contribute to myocardial remodeling through altered ECM organization, inflammatory communication, and crosstalk with cardiomyocytes, resident immune cells, and endothelial cells. Green indicates upregulated genes and red indicates downregulated genes in HOCM-CFs.

### Study limitations

The number of control CFs was limited to three samples and were not ethnicity-matched to HOCM samples, which may influence transcriptomic comparisons. The comparison with myocardial tissue RNA-seq was exploratory, and was limited by tissue heterogeneity, sample size, and lack of cell-type resolution. Future studies using single-cell or spatial transcriptomics, larger cohorts, and functional experiments will be required to define the cellular origin, spatial localization, and mechanistic role of the identified inflammatory and ECM-remodeling pathways.

## Data availability statement

The datasets generated and analysed in the current study are available in the GEO repository: GEO (GSE337273).

## Author Contributions

AMI, YA, and MY conceived and designed the study. AG performed the RNA-sequencing experimental workflow. SH, AG, AMI, and LRS optimized the RNA-sequencing analysis pipeline and performed differential expression, visualization, network, and functional enrichment analyses. MR and AK contributed to the interpretation of the sequencing data. AMI, SE, HE, MO, FM, and MR contributed to sample collection and performed cardiac fibroblast experiments, including cell isolation, characterization, RNA/protein validation, and validation data analysis. MH and MY performed patient clinical profiling, data curation, and assessment of clinical relevance. MA Performed cardiovascular-genetic examination and analysis of HOCM patients. All authors reviewed and approved the final version of the manuscript.

## Nonstandard Abbreviations and Acronyms

BNP: B-type natriuretic peptide
CFs: cardiac fibroblasts
CMR: cardiac magnetic resonance
CMs: cardiomyocytes
COL: collagen
CXCR4: C-X-C chemokine receptor 4
ECM: extracellular matrix
HOCM: hypertrophic obstructive cardiomyopathy
LVOTO: left ventricular outflow tract obstruction
MMP: matrix metalloproteinase
TGFβ1: transforming growth factor beta 1

